# Identification of SSX2IP/Msd1 as a Wtip-binding partner by targeted proximity biotinylation

**DOI:** 10.1101/2021.05.16.444380

**Authors:** Bo Xiang, Alice H. Reis, Keiji Itoh, Sergei Y. Sokol

## Abstract

Wilms tumor-1-interacting protein (Wtip) is a LIM-domain-containing adaptor that links cell junctions with actomyosin complexes and modulates actomyosin contractility and ciliogenesis in *Xenopus* embryos. The Wtip C-terminus with three LIM domains binds binds Shroom3 and modulates Shroom3-induced apical constriction in ectoderm cells. We found that the N-terminal domain localizes to the basal bodies in skin multiciliated cells, but its interacting partners remain largely unknown. Using a novel targeted proximity biotinylation approach with anti-GFP antibody attached to the biotin ligase BirA in the presence of GFP-Wtip-N, we identified SSX2IP as the candidate binding protein. SSX2IP, also known as Msd1 or ADIP, is a centriolar satellite protein that functions as a targeting factor for ciliary membrane proteins. Wtip physically associated with SSX2IP and the two proteins formed mixed spherical aggregates in overexpressing cells in a dose-dependent manner, in a process that resembles phase separation. These results suggest that the interaction between SSX2IP and Wtip is relevant to their functions at the centrosome and basal bodies. The described antibody targeting of biotin ligase should be applicable to other GFP-tagged proteins.

## INTRODUCTION

The Ajuba family of LIM-domain-containing proteins are adaptors that participate in various processes that involve actomyosin contractility. Ajuba localizes to adherens junctions and modulates Rac1 activity in mammalian cells (Kadrmas and Beckerle, 2004; Marie et al., 2003; Pratt et al., 2005; Schimizzi and Longmore, 2015). At the junctions, Ajuba associates with α-catenin and inhibits Hippo signaling in a tension-dependent manner (Das Thakur et al., 2010; Nola et al., 2011; Rauskolb et al., 2014). Also, Ajuba homologs localize to the centrosomes of mammalian and Drosophila cells and function in cell division and mitotic spindle orientation (Hirota et al., 2003; Sabino et al., 2011; Schimizzi and Longmore, 2015).

Wilms tumor 1-interacting protein (Wtip) is one of Ajuba homologues. Wtip is composed of two conserved domains that exhibit distinct properties. The C-terminus of Wtip contains three LIM domains and interacts with Shroom3, an actin binding protein that can ectopically induce apical constriction in different cell types (Haigo et al., 2003; Hildebrand and Soriano, 1999). Wtip also binds the core planar cell polarity (PCP) protein Prickle3 and has been implicated in PCP and ciliogenesis in *Xenopus* and zebrafish embryos (Bubenshchikova et al., 2012; Chu et al., 2016; Chu et al., 2018). Furthermore, the C-terminal domain of Wtip has been found in the nucleus, where it is proposed to associate with transcriptional repressors of the Snail family and regulate transcription (Langer et al., 2008). By contrast, the functions of the Wtip N-terminal domain that includes the first 480 amino acids of the protein (WtipN) are largely unknown. We find that the WtipN localizes to puncta at ectodermal cell junctions and to basal bodies in the epidermal multiciliated cells (Chu et al., 2018), however, the molecular mechanisms underlying its functions remain to be elucidated.

In this study, we searched for proteins associating with WtipN using a novel proximity biotinylation approach. The promiscuous bacterial biotin ligase BirA* has been fused with the GFP-specific single domain antibody, also known as GFP-binding protein (GBP)(Kubala et al., 2010; Rothbauer et al., 2008). This fusion protein (BirA-GBP) would be expected to target the biotinylation activity to the immediate proximity of GFP-WtipN in vivo. This approach identified SSX2IP, also known as Msd1 and ADIP (Asada et al., 2003)) as one of the main candidates. Interestingly, SSX2IP is a maturation factor for mitotic centrosomes (Barenz et al., 2013), and a targeting factor for ciliary membrane proteins cooperating with Cep290, the BBSome, and Rab8 (Klinger et al., 2014). SSX2IP also interacts with Afadin and a-Actinin (Asada et al., 2003) and is implicated in linking cell junction to actin cytoskeleton. This interaction of Wtip and SSX2IP has been further validated by the immunoprecipitation and the colocalization of the two proteins in ectoderm cells. Thus, the targeted proximity biotinylation (TPB) approach identified a new interaction partner for Wtip and will likely be useful for defining protein-protein interactions in studies of other GFP-tagged proteins.

## MATERIALS AND METHODS

### Xenopus embryo culture and microinjections

In vitro fertilization and culture of Xenopus laevis embryos were carried out as previously described (Dollar et al., 2005). Animal care and use was in accordance with the NIH guidelines as established by the Icahn School of Medicine at Mount Sinai, New York. Staging was according to Nieuwkoop and Faber (Nieuwkoop and Faber, 1967). For microinjections, four-cell embryos were transferred into 3 % Ficoll in 0.5 x MMR buffer and 10 nl of mRNAs with or without biotin solution was injected into one or more blastomeres. Amounts of injected mRNA or biotin per embryo have been optimized in preliminary dose-response experiments and are indicated in figure legends.

### Plasmids, mRNA synthesis

WtipN encodes the N-terminal 480 amino acids of Xenopus Wtip. The plasmids for *Xenopus* Wtip including Flag-Wtip, GFP-WtipN, GFP-WtipC, RFP-WtipN, Flag-WtipN, Flag-GFP, GFP-Prickle3C and Flag-Prickle3C have been previously described (Chu et al., 2016; Chu et al., 2018). A plasmid encoding a mutated myc-BirA (Roux et al., 2012), a gift from Brian Burke, was subcloned into pCS2 vector. WtipN was fused to BirA by subcloning in pCS2-myc-BirA vector to generate WtipN-MycBirA. BirA DNA was fused to the N-terminus of the DNA fragment encoding a single domain antibody specific for GFP (Kubala et al., 2010) in pCS2 vector (BirA-GBP). The insert from the human Msd1/SSX2IP plasmid (a gift from T. Toda) was subcloned into pCS2-GFP vector to make GFP-hSSX2IP. *Xenopus* SSX2IP cDNA clone (SSX2IP.S) was obtained from Dharmacon and the insert was subcloned into pXT7-EGFP to make GFP-SSX2IP-pXT7. Details of cloning are available upon request. Capped mRNAs were made by in vitro transcription from linearized plasmids as templates with T7 or SP6 RNA polymerases using mMessage mMachine kit (Ambion).

### Immunoblot analysis

Immunoblotting was carried out as previously described (Itoh et al., 2005). Briefly, whole embryos were lysed in a buffer containing 1 % Triton X-100, 50 mM sodium chloride, 50 mM Tris-HCl at pH 7.6, 1 mM EDTA, 0.6 mM phenylmethylsulphonyl fluoride (PMSF), 10 mM sodium fluoride and 1 mM sodium orthovanadate. The lysates were cleared and subjected to SDS-polyacrylamide gel electrophoresis, and proteins were transferred to the PVDF membrane for immunodetection with the indicated antibody. Antibodies against the following antigens were used: myc (1:200, mouse monoclonal 9E10 hybridoma supernatant), His (1:1000, Invitrogen, mouse monoclonal), Flag M2(1:1000, mouse monoclonal, Sigma), GFP (1:1000, B-2, Santa Cruz, mouse monoclonal), Biotin (1:10000, goat polyclonal, Pierce). Secondary antibodies were against mouse or goat IgG conjugated to HRP (1:3000, Jackson Immunoresearch or Santa Cruz). Biotinylated proteins were detected similarly with the following modifications. Membranes were blocked in 3 % nonfat milk in PBS with 0.1% Tween 20 and incubated in the same buffer with HRP-conjugated streptavidin (1:5,000; Invitrogen).

For co-immunoprecipitation of Flag tagged Wtip or GFP constructs with GFP-SSX2IP, 30 embryos at 4-8 cell stages were injected with 330 pg Flag-Wtip, Flag-WtipN or Flag-GFP and 300 pg GFP-SSX2IP RNA into four animal blastomeres. When control embryos reached stage 11, embryos were lysed in 500 µl of the lysis buffer and the lysates were cleared by centrifugation at 16,000 g for 4 min to remove yolk platelets. The supernatant was incubated with 2 µl of anti-Flag M2 agarose beads overnight at 4 degrees. The beads were washed 5 times with the lysis buffer. 20 µl of 2 x SDS sample buffer were added to the beads and boiled for 5 min. 15 ml of the sample were analyzed in Western blotting.

### Affinity capture of biotinylated proteins

100 embryos at 4-8 cell stages were injected four times into animal blastomeres with 10 nl containing RNAs and Biotin (1.2 mM) and cultured until st. 12.5 or st. 14. Then, the embryos were lysed in 1 ml of the lysis buffer and the lysates were centrifuged at 16,000 g for 4 min. Supernatants were incubated with 40 μl avidin beads (Invitrogen) at 4°C overnight. The beads were washed with PBS four times. Bound proteins were collected from the agarose beads with 50 μl of the SDS-sample buffer at 98°C for 5 min. 10 % of the sample was reserved for Western blot analysis. For the larger scale preparation, 90% of the sample were loaded on a gel.

### Protein identification by mass spectrometry

Proteins eluted from the avidin beads by boiling in the SDS containing sample buffer and separated by SDS-PAGE using standard procedures (Itoh et al., 2005). Separated proteins were visualized by Coomassie Blue staining. The whole gel lanes were cut in several gel bands and submitted to in-gel trypsin digestion and LC-MS/MS analysis at the Keck Proteomics Laboratory (Yale University). Data were annotated using *Xenopus* genome version 9.2 at www.xenbase.com. Results were analyzed by the Scaffold software. The acceptance level for proteins was two identified peptides with minimum 95% probability each.

## Results

### WtipN is localized to the basal bodies of multiciliated skin cells

We have previously described the subcellular distribution of Wtip at junctional puncta in epidermal ectoderm and neuroectoderm at late gastrula/early neurula stages (Chu et al., 2018) and it was found at the centrosomes of the ciliated endoderm at the *Xenopus* gastrocoel roof (Chu et al., 2016). To further explore the localization of WtipN, encoding the first 480 amino acids of *Xenopus* Wtip, we microinjected GFP-WtipN and GFP-WtipC RNAs and monitored the embryos at the beginning of gastrulation and at the tailbud. Consistent with our previous observations, GFP-WtipN was detected at cell junctions, whereas GFP-WtipC was in the nuclei in early gastrula ectoderm (**Fig. 1A, B**). Notably, at tailbud stages, GFP-Wtip localized to basal bodies of skin multiciliated cells as well as cell junctions (**Fig. 1C**). These results further confirm the association of WtipN with the centrosome/basal body. Since the centrosomal proteins interacting with WtipN have not been reported, with the exception of Prickle3 (Chu et al., 2016), a search for such proteins is warranted.

**Figure 1.**
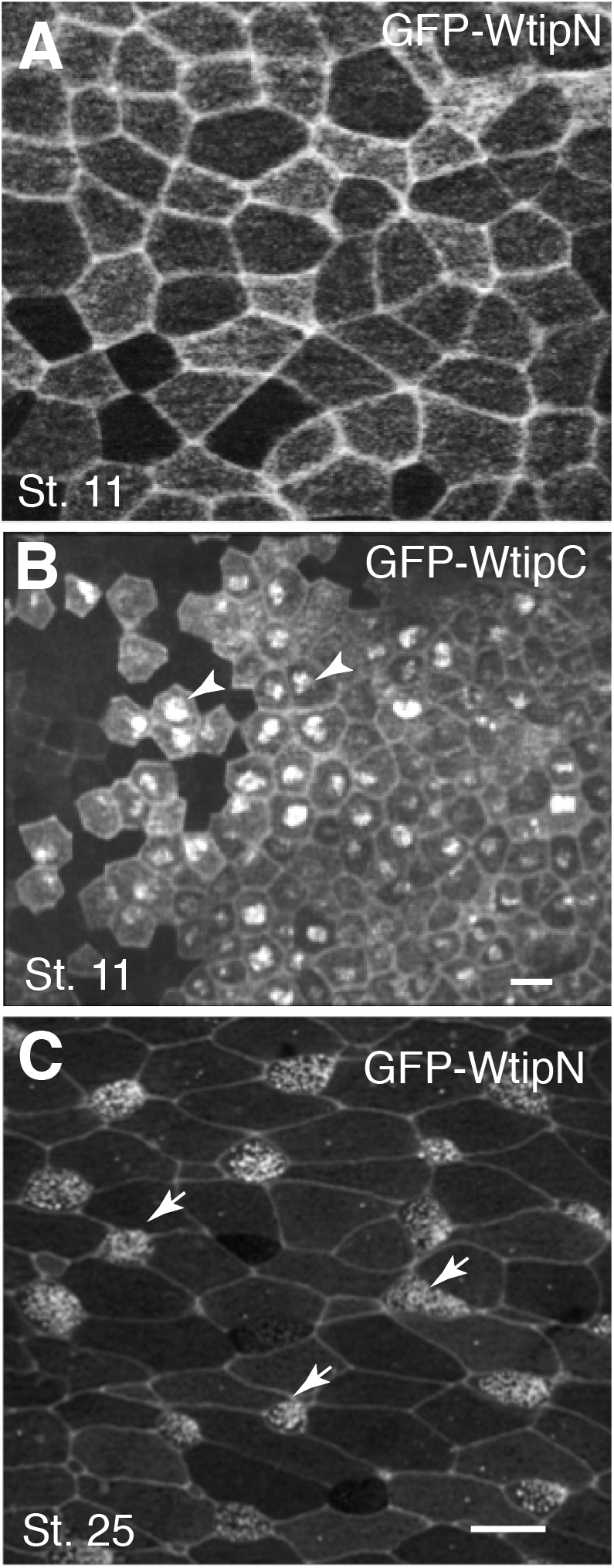
Localization of WtipN constructs in basal bodies of multiciliated cells. Embryos were injected with 200 pg of GFP-WtipN (A, C) or GFP-WtipC RNA (B), fixed and imaged at the indicated stages. A, GFP-WtipN junctional staining in stage 10.5 ectoderm. B, WtipC is predominantly in the nuclei of the ectoderm cells (stage 10.5). C, GFP-WtipN is localized to basal bodies of skin multiciliated cells (arrows, stage 25). Scale bars in B (also applies to A) and C are 20 µm.

### Biotinylation of Wtip by BioID fusions in *Xenopus* embryos

To search for new interacting partners of Wtip, we decided to use proximity biotinylation or BioID (Roux et al., 2012) that allows to isolate associated proteins by labeling them *in vivo* with biotin using a mutated bacterial biotin ligase. We generated a fusion of BirA with WtipN (myc-BirA-WtipN) and tested its enzymatic activity in vivo, in Xenopus embryos. To optimize the *in vivo* BioID protocol, RNAs for myc-BirA or myc-BirA-WtipN were coinjected with or without biotin into four-cell embryos and cultured the embryos until late gastrula or early neurula stages. We observed robust self-biotinylation of both myc-BirA-WtipN and myc-BirA in the presence of biotin starting as early as stage 10.5 (**Fig. 2A** and not shown). Notably, in the absence of biotin, the background was low, except for two major protein bands around 70 and 120 kDa recognized by Streptavidin-HRP, possibly representing endogenous biotin-containing proteins. Pulldowns with Streptavidin beads contained many bands in the sample from myc-BirA-WtipN-expressing embryos (**Fig. 2B**), which likely correspond to WtipN-interacting proteins.

**Figure 2.**
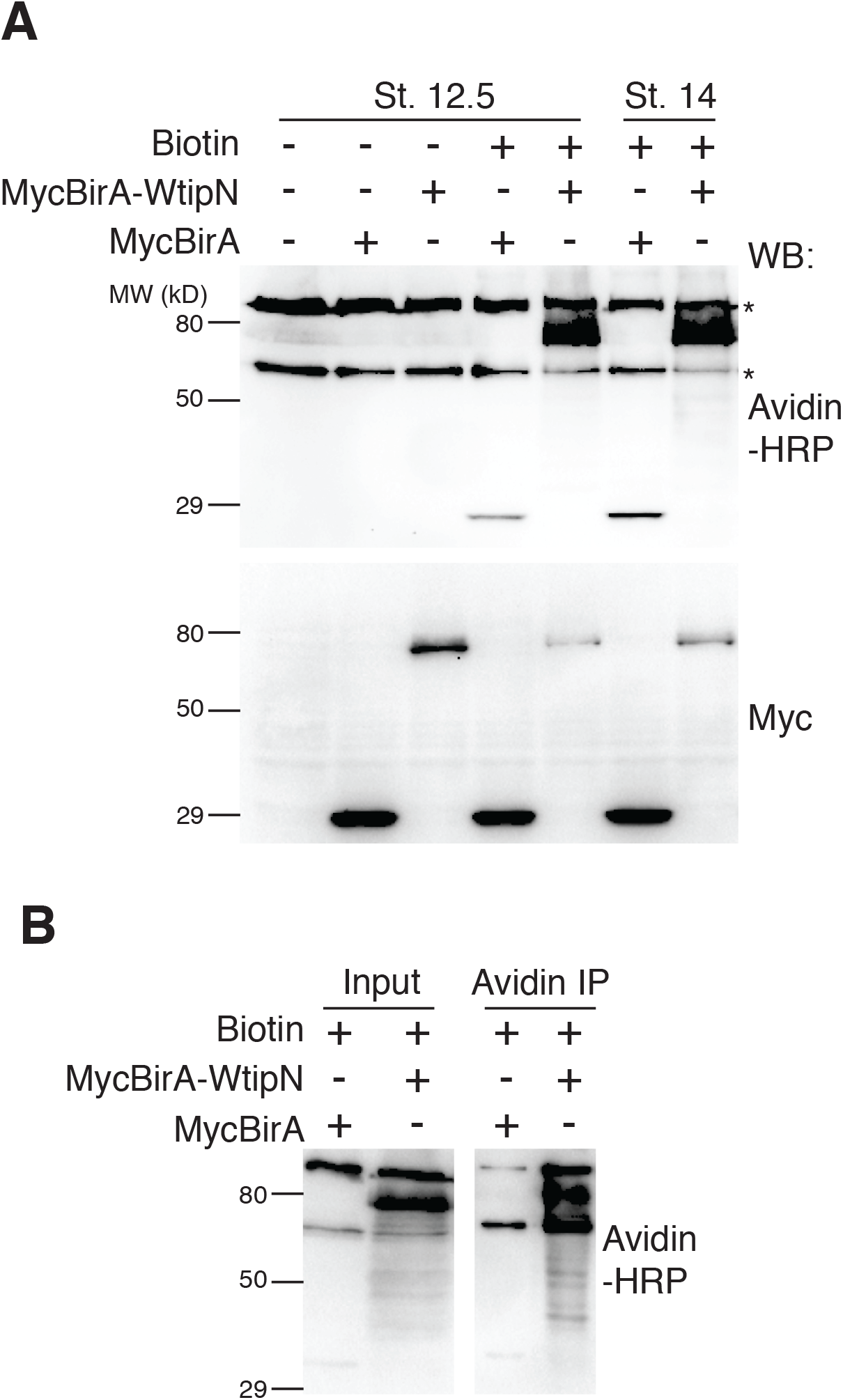
Biotinylation of the BirA-WtipN fusion in Xenopus embryos. Four-cell embryos were injected with 300 pg of myc-BirA RNA or 720 pg of mycBirA-WtipN RNA with or without 0.8 mM biotin. A. Immunoblotting with Avidin-HRP reveals self-biotinylation of mycBirA and mycBirA-WtipN only in the presence of biotin. Protein levels were assessed with anti-myc antibody. B. Biotinylated proteins were pulled down with avidin beads and visualized by Avidin-HRP. In addition to self-biotinylated mycBirA-Wtip, multiple biotinylated bands are detected in the lysates as well as in pulldown.

### Targeted proximity biotinylation

We next modified the approach by fusing BirA to the coding sequence of a single domain antibody against GFP, or GFP-binding protein (Kubala et al., 2010) (BirA-GBP). Such fusion would be predicted to biotinylate any GFP-tagged protein and others in its immediate proximity. This construct can serve as a universal tool for <u>targeted proximity biotinylation</u> (TPB). To validate the efficiency and specificity of this approach, we injected BirA-GBP RNA into early embryos together with GFP-WtipN and Flag-WtipN.

As expected, we observed an efficient and specific biotinylation of GFP-WtipN but not Flag-WtipN, when it was coexpressed with BirA-GBP and biotin (**Fig. 3A**). Of note, BirA-GBP failed to biotinylate FlagGFP, possibly due to lack of suitable lysine sites (**Fig. 3A**). To show that TPB can be applied to a different protein, we confirmed that BirA-GBP biotinylates GFP-Prickle3C but not Flag-Prickle3C (Chu et al., 2016) **(Fig. 3B)**. These results suggest that TPB is applicable to protein network analysis for any GFP fusion protein.

**Figure 3.**
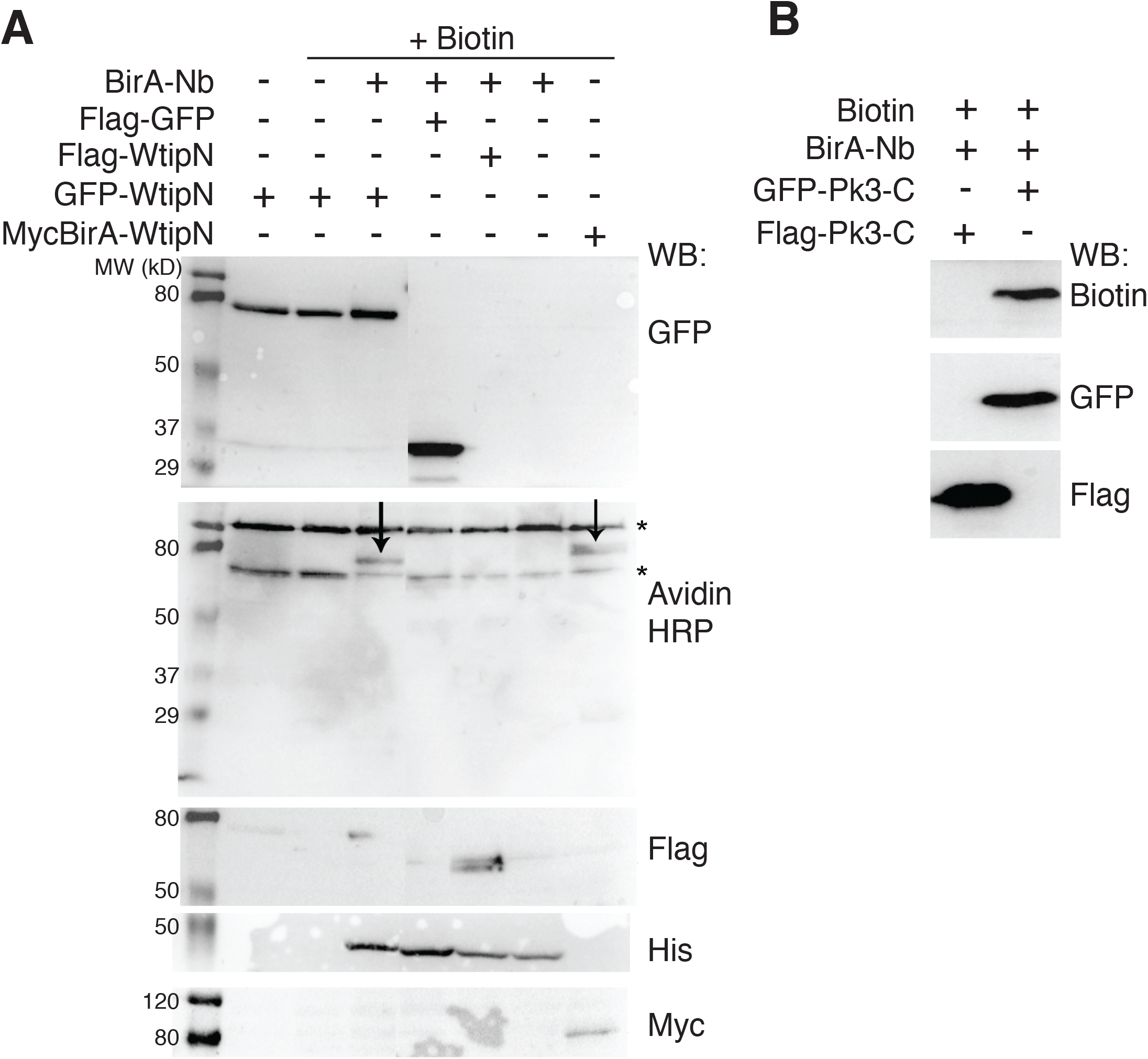
Specificity of BirA-GBP-mediated biotinylation. A, GFP-WtipN but not Flag-WtipN is biotinylated by BirA-GBP in the presence of biotin. MycBirA-WtipN is a positive control for biotinylation. B, GFP-Prickle3C protein, but not Flag-Prickle3C, was biotinylated by BirA-GBP. Protein expression levels are assessed with tag-specific antibodies as indicated (A, B).

### Identification of SSX2IP/Msd1/ADIP as a novel interacting partner of WtipN

To search for proteins interacting with WtipN, we carried out proximity biotinylation reaction in vivo, in embryos coexpressing GFP-WtipN and BirA-GBP in the presence of biotin. Embryos coexpressing Flag-WtipN and BirA-GBP were used as negative controls (**Fig. 4A**). A separate group of embryos was injected with Myc-BirA-WtipN. Biotinylated proteins were captured with neutravidin-agarose beads, the bound proteins were separated by SDS-PAGE separation and visualized by Simple Blue staining. Gel slices were analyzed by LC-MS/MS to identify candidate interacting proteins. This analysis identified several proteins that were present in the group of GFP-WtipN and BirA-GBP, but absent in the group with Flag-WtipN and BirA-GBP. Multiple identified peptides belonged to the *Xenopus* homologue of SSX2IP/ADIP/Msd1 (Asada et al., 2003; Barenz et al., 2013; Toya et al., 2007). The same protein was identified at high confidence using Myc-BirA-WtipN as a bait (**Fig. 4B**). These findings indicate that SSX2IP and WtipN interact.

**Figure 4.**
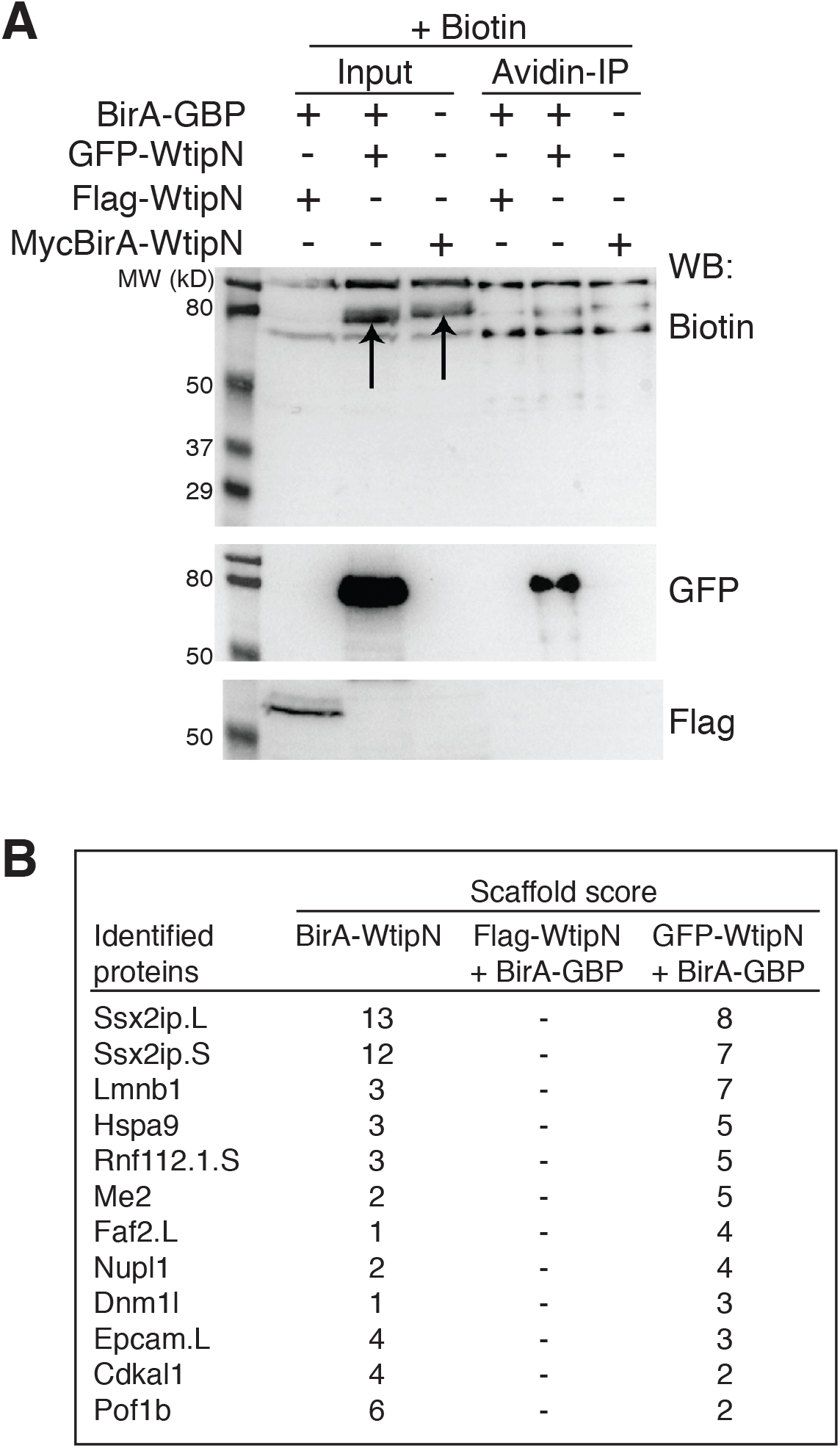
Identification of SSX2IP by targeted proximity biotinylation. Early embryos were injected with indicated RNAs and biotin (1.2 mM). A. Biotinylated protein were enriched with avidin agarose beads. 1.5 μL elution from 30 μL were loaded and visualized with indicated antibodies. Biotinylated GFP-WtipN but not Flag-WtipN in the presence of BirA-GBP was detected in the pulldown. MycBirA-WtipN was biotinylated in the absence of BirA-GBP. B. Top hits identified by LC-MS/MS analysis that are present in the experimental groups, but not in the negative control group.

### SSX2IP associates with WtipN in embryonic ectoderm

We next validated the identified interaction by co-immunoprecipitation. Flag-WtipN protein strongly associated with GFP-SSX2IP protein at the beginning of gastrulation when co-expressed in *Xenopus* embryos. However, the full length Wtip protein was less efficient in the binding of GFP-SSX2IP protein (**Fig. 5)**. These experiments strongly suggest that SSX2IP might associate with WtipN in vivo. Of note, both Wtip and SSX2IP are found at or near the basal bodies (Chu et al., 2016; Chu et al., 2018; Hori et al., 2014; Klinger et al., 2014).

**Figure 5.**
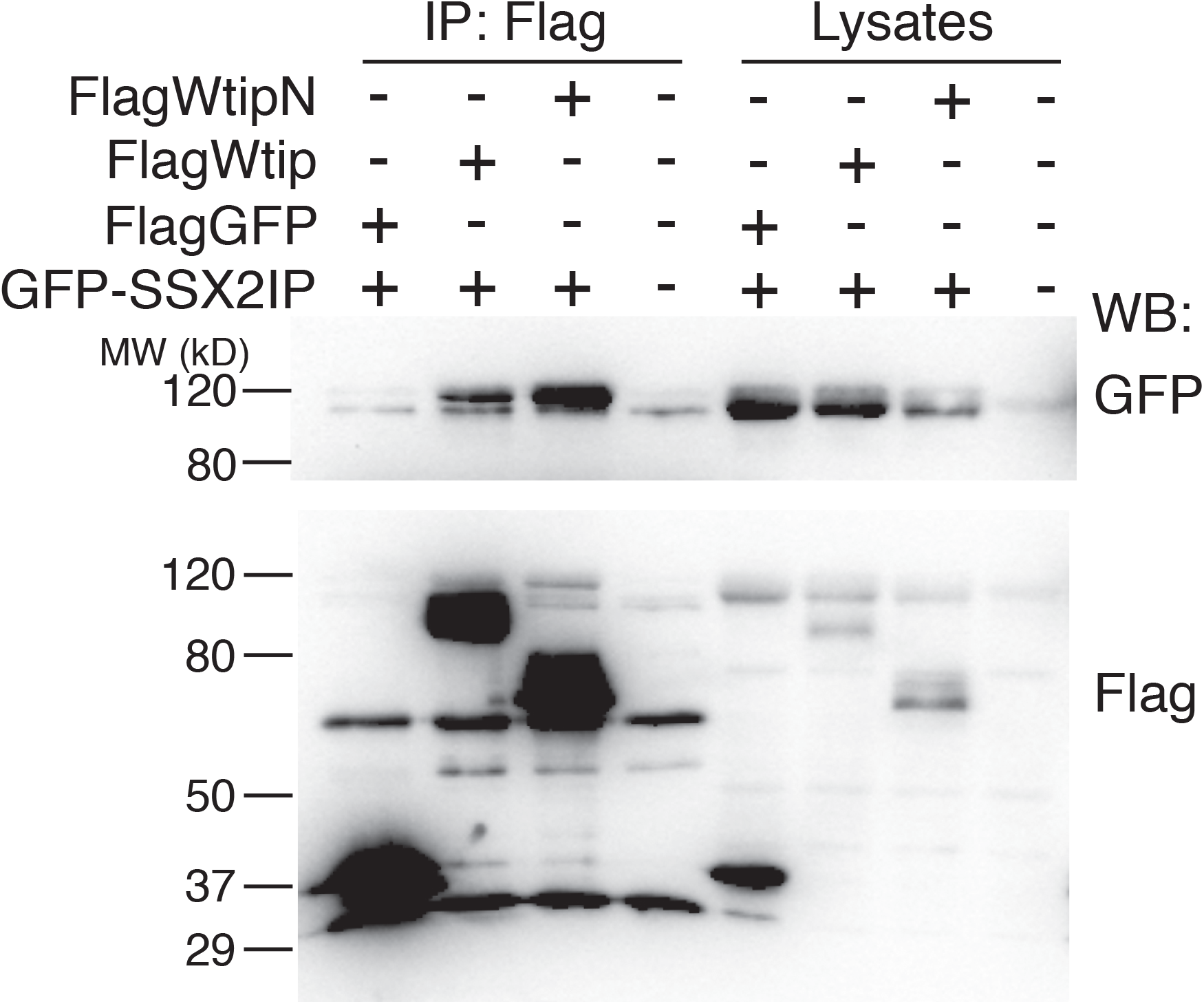
The association of ADIP and Wtip. A. Four-cell embryos were coinjected with GFP-SSX2IP RNA (300 pg) and either Flag-WtipN or Flag-Wtip RNA (330 pg). Flag-GFP RNA (330 pg) was the negative control. Embryos were lysed at stage 11 and lysates were immunoprecipitated with anti-Flag agarose beads and immunoblotted with antibodies specific for Flag and GFP. WtipN was more strongly co-immunoprecipitated with ADIP than wild type Wtip.

We next tested whether SSX2IP colocalizes with WtipN in ectodermal cells. When two RNAs were expressed in early embryos, GFP-SSX2IP and Flag-WtipN revealed striking colocalization in cytoplasmic aggregates in early ectoderm (**Fig. 6**). The SSX2IP/WtipN protein complexes were of spherical shape and exhibited dose-dependent size increase, suggesting an aggregation process that is similar to liquid phase separation (Alberti and Hyman, 2021). Together these experiments suggest that SSX2IP binds the N-terminal fragment of Wtip and likely mediates its function at the same subcellular locations.

**Figure 6.**
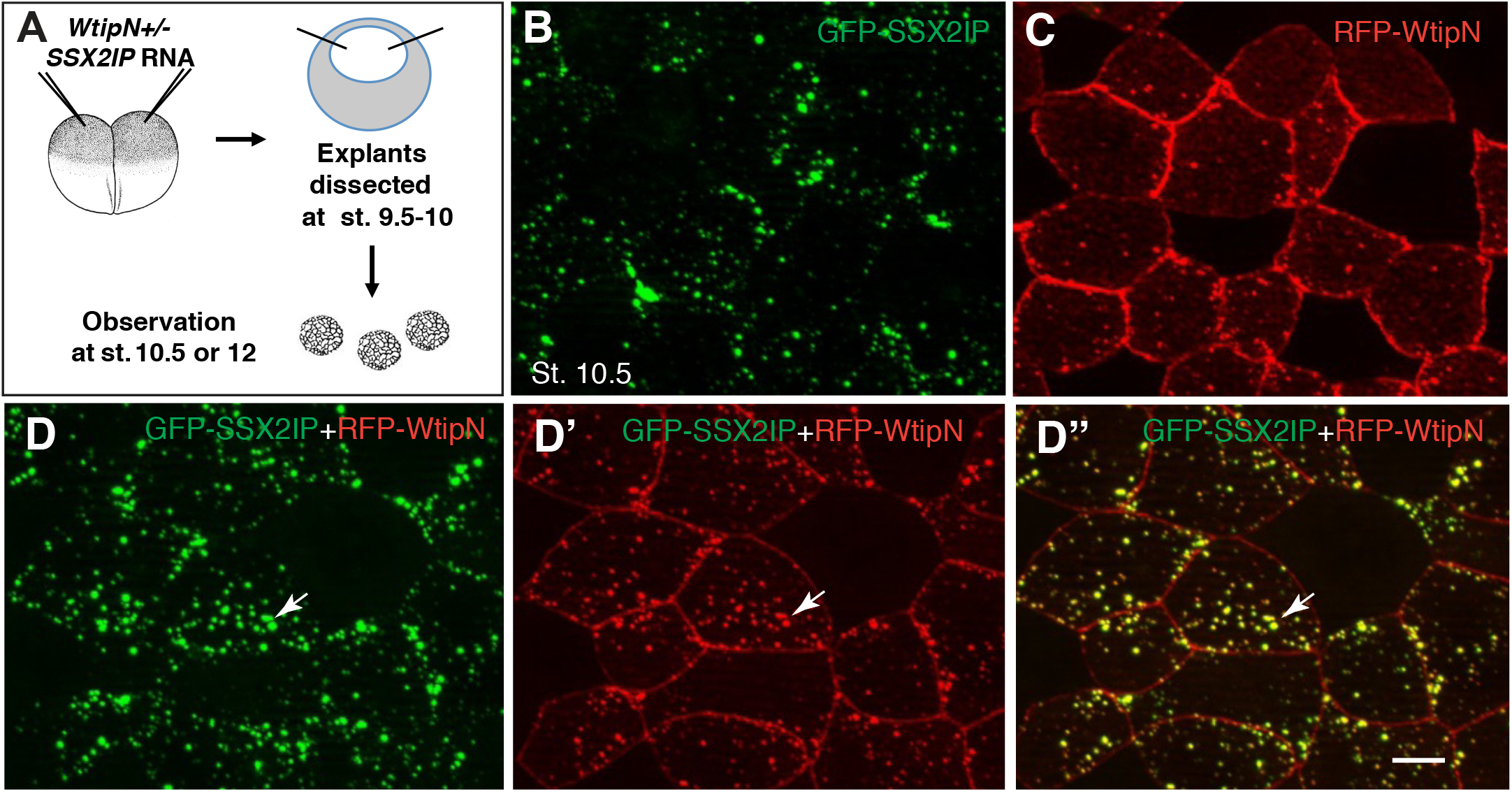
SSX2IP colocalizes with WtipN in vivo. RFP-WtipN RNA (300 pg) was coinjected with GFP-SSX2IP (300 pg) at 4-cell embryos. When the embryos reached stage 12.5, they were fixed, and the ectodermal tissue was imaged. SSX2IP colocalizes with WtipN in cytoplasmic puncta.

## Discussion

In this study, we used proximity biotinylation to search for new interaction partners of Wtip, a LIM-domain protein of the Ajuba/Zyxin family. Wtip is present in many tissues and has been shown to localize to the centrosome/basal bodies and cell junctions (Chu et al., 2016; Chu et al., 2018). This diverse localization may underlie the proposed functions of Wtip in morphogenesis and planar cell polarity as well as its roles in the centrosomal organization and ciliogenesis (Bubenshchikova et al., 2012; Chu et al., 2016; Chu et al., 2018). Wtip has been also implicated in forming a complex with the Snail transcription factors leading to the activation of neural crest markers (Langer et al., 2008). The knowledge of new interaction partners of Wtip should help elucidate how it functions in the cell.

The BioID (biotin identification) approach is useful for the detection of protein interactions under physiological conditions, overcoming non-specific contamination that often accompanies traditional pulldowns. This approach is more sensitive and specific as compared to conventional immunoprecipitation or yeast-two-hybrid screens, because it can detect weak, transient, or hydrophobic protein interactions in their native state (Roux et al., 2018). We modified this technique by fusing BirA with an anti-GFP antibody that brings the enzyme into close proximity with GFP-WtipN. One advantage of this approach is that the generation of the direct fusion of the ‘bait’ protein with BirA is no longer needed. Also, this method may identify a more diverse protein population due to the increased distance between the enzyme and the bait. Importantly, we identified SSX2IP using both BirA-GBP/GFP-WtipN and WtipN-BirA, increasing our confidence in the result. Thus, our findings and a similar report using zebrafish embryos (Xiong et al., 2021) indicate that TPB represents a versatile approach for protein interaction studies of various GFP-tagged proteins.

Our analysis identified SSX2IP/ADIP/Msd1 as a protein associating with Wtip N-terminus. SSX2IP is a highly conserved centriolar satellite protein that functions in ciliogenesis (Hori et al., 2014) and anchoring cytoplasmic microtubules to the centrosome (Toya et al., 2007). We validated the binding using standard immunoprecipitation and *in vivo* colocalization of the two proteins. The implication of both Wtip and SSX2IP in ciliogenesis (Bubenshchikova et al., 2012; Hori et al., 2014) suggests a potential role of this interaction in PCP signaling. Notably, spherical aggregates composed of both WtipN and SSX2IP formed in the cytoplasm, and their size was dependent on the dose of injected RNAs. Such aggregation process is similar to known cases of phase separation that has been observed for the centrosome assembly both *in vitro* and *in vivo* (Alberti and Hyman, 2021; Wheeler and Hyman, 2018; Woodruff et al., 2015). Whether the identified WtipN-SSX2IP interaction contributes to the assembly of the pericentriolar material or has a different physiological significance remains to be clarified in future studies.

Like SSX2IP, Wtip is also present at the centrosome and basal bodies of different cell types and appears to require its N-terminus for this localization (Chu et al., 2016; Chu et al., 2018). Thus, the interaction of Wtip with SSX2IP may be a prerequisite for the centrosomal/basal body localization for one or the other protein. Besides the roles at the centrosome and/or basal body, the association of Wtip and SSX2IP may play a role in actomyosin dynamics. Both proteins interact with tension-sensitive molecules that organize actomyosin complexes at cell junctions (Asada et al., 2003; Chu et al., 2018). Further analysis is needed to demonstrate whether SSX2IP contributes to regulation of actomyosin contractility.

## Acknowledgements

We thank Olga Ossipova for her comments and help with the writing of this manuscript. We also thank Andriani Ioannou and Fei Wu who contributed to the early stages of this work. This study has been supported by the NIH grant R35GM122492 to SYS.

